# 3-nitrotyrosine shortens axons of a non-dopaminergic neuron by inhibiting mitochondrial motility

**DOI:** 10.1101/2024.05.09.593299

**Authors:** Masahiro Hirai, Kohei Suzuki, Yusuke Kassai, Yoshiyuki Konishi

**Author notes:** Authors contributed equally (Masahiro Hirai, Kohei Suzuki). **Corresponding author** Yoshiyuki Konishi, Department of Applied Chemistry and Biotechnology, Faculty of Engineering, University of Fukui, 3-9-1 Bunkyo, Fukui, Fukui 910-8507, Japan.

## Abstract

3-nitrotyrosine (3-NT), a byproduct of oxidative/nitrosative stress, is implicated in age-related neurodegenerative disorders. Existing literature indicates that free 3-NT becomes integrated into the carboxy-terminal domain of α-tubulin through the tyrosination/detyrosination cycle. Independently of this integration, 3-NT has been linked to the cell death of dopaminergic neurons. Given the critical role of tyrosination/detyrosination in governing axonal morphology and function, the substitution of tyrosine with 3-NT in this process may potentially disrupt axonal homeostasis, although this aspect remains underexplored. In this study, we examined the impact of 3-NT on the axons of cerebellar granule neurons, representing non-dopaminergic neurons. Our observations revealed axonal shortening, which correlated with the incorporation of 3-NT into α-tubulin. Importantly, this axonal effect was observed prior to the onset of cellular death. Furthermore, 3-NT was found to diminish mitochondrial motility within the axon, resulting in a subsequent reduction in mitochondrial membrane potential. The suppression of syntaphilin, a protein responsible for anchoring mitochondria to microtubules, reversed the 3-NT-induced decrease in mitochondrial motility, consequently restoring axonal elongation. These findings underscore the inhibitory role of 3-NT in axonal elongation by impeding mitochondrial movement, implying its potential involvement in axonal dysfunction within non-dopaminergic neurons.

## 1. Introduction

The aging brain undergoes structural and functional changes accompanied by cognitive decline and volumetric loss (Blinkouskaya et al., 2021). Emerging evidence suggests that neuronal atrophy, encompassing axons, dendrites, and synapses, occurs during normal brain aging and in age-associated neurodegenerative disorders preceding widespread cell death (Hof and Morrison, 2004; Dickstein et al., 2013; Stahon et al., 2016; Lee and Kim, 2022). However, the precise mechanisms underlying the effects of aging on neuronal morphology remain to be fully elucidated. Reactive oxygen species play pivotal roles in both the aging process and neurodegenerative disorders (Lin and Beal, 2006; Niedzielska et al., 2016). Superoxide, upon reacting with nitric oxide, generates peroxynitrite, which in turn leads to the formation of 3-nitrotyrosine (3-NT) through interaction with protein tyrosine residues or free tyrosine (Drew et al., 2002; Radi, 2004, 2018; Pacher et al., 2007; Souza et al., 2008). Notably, 3-NT has garnered attention not only as a marker of oxidative stress but also as a mediator of cell death and age-related disorders (Tohgi et al., 1999; Butterfield et al., 2007; Larsen et al., 2008; Andreazza et al., 2009; Franco and Estévez, 2014; Bandookwala and Sengupta, 2020). Previous investigations have suggested that free 3-NT selectively induces toxicity in dopaminergic (DA) neurons via the activity of aromatic amino acid decarboxylase, potentially implicating its role in the pathogenesis of Parkinson’s disease (Mihm et al., 2001; Blanchard-Fillion et al., 2006).

α-tubulins undergo cycles of addition and removal of a tyrosine residue at the carboxy-terminal, regulated by tubulin tyrosine ligase (Ttl) (Ersfeld et al., 1993; Erck et al., 2005), vasohibins, or microtubule-associated tyrosine carboxypeptidase (Aillaud et al., 2017; Nieuwenhuis et al., 2017; Landskron et al., 2022). Previous studies have shown that 3-NT is selectively incorporated into α-tubulins instead of tyrosine, albeit without apparent involvement in neuronal cell death (Eiserich et al., 1999; Kalisz et al., 2000; Pellufo et al., 2004; Blanchard-Fillion et al., 2006). The ratio of tyrosination to detyrosination of α-tubulin varies between axons and dendrites, as well as among axonal branches (Witte et al., 2008; Seno et al, 2016; Imanaka et al., 2022), and is believed to exert significant control over axonal morphology (Erck et al., 2005; Konishi and Setou, 2009; Aillaud et al., 2017). Disruption of tyrosination or detyrosination of α-tubulin alters kinesin- or dynein-mediated axonal transport and the function of microtubule-associated proteins (Erck et al., 2005; Konishi and Setou, 2009; Peris et al., 2009; McKenney et al., 2016; Nirschl et al., 2016) resulting in defects in axonal function and morphology. Hence, the incorporation of 3-NT into α-tubulin may disrupt these processes across different neuron types. Indeed, a recent study demonstrated that exposure to agrochemicals inhibits axonal transport of mitochondria in dopaminergic neurons carrying mutations associated with Parkinson’s disease in the α-synuclein gene, possibly mediated by α-tubulin nitration (Stykel et al., 2018). However, whether 3-NT disrupts the regulation of axonal morphology in both dopaminergic and non-dopaminergic neurons remains to be determined.

In this study, we aimed to elucidate the potential consequences of 3-NT on neuronal axons in general, utilizing cerebellar granule neurons (CGNs), which are non-dopaminergic. Our findings reveal that 3-NT impedes axonal elongation by arresting mitochondrial axonal transport, offering a potential mechanistic link between oxidative stress and disruption of axonal structure.

## 2. Methods

### 2.1. Primary Culture of CGNs

Animal handling procedures were conducted under controlled environmental conditions, with temperatures maintained at 18–28°C and a regulated light cycle of 10 hours light and 14 hours dark. Animals had access to food and water ad libitum. CGNs were prepared from Jcl:ICR mice (CLEA Japan, Tokyo, Japan) at postnatal days 4 to 6 as described previously (Kubota et al., 2013, Matsumoto et al., 2022). Following decapitation, isolated cerebellar were digested with trypsin (10 mg/ml)-DNase (200 units/ml) (TRL, DP, Worthington, Lakewood, NJ) solution. The dissociated CGNs were suspended in Minimal Essential Medium (11090-081, Thermo Fisher Scientific, Waltham, MA) supplemented with 10% calf serum (SH30072.03, Thermo Fisher Scientific), penicillin (100 units/ml, P7794, Sigma-Aldrich, St. Louis, MO), streptomycin (0.1 mg/ml, P9137, Sigma-Aldrich), glutamine (2 mM, G8540, Sigma-Aldrich), and KCl (25 mM, 163-03545, FUJIFILM Wako Pure Chemical Corporation, Osaka, Japan). For live-cell imaging of CGNs (as depicted in Fig. 4–6), Minimal Essential Medium without phenol red (51200–038, Thermo Fisher Scientific) supplemented with 20 mM Hepes-KOH (340-01371, Dojindo Laboratories, Kamimashiki-gun, Japan, 168-21815, FUJIFILM Wako Pure Chemical Corporation) at pH 7.2 and other required supplements was employed as the culture media.

In the experiments illustrated in Fig. 1, Fig. 2, and Fig. 3A, CGNs were seeded in 12-well plates with cover glass (C018001, Matsunami Glass, Osaka, Japan) previously coated with 15 μg/ml poly-L-ornithine (P2533, Sigma-Aldrich) at a density ranging from 0.5 to 1 × 10^6^ cells/well, depending on the specific experiment. For the experiments depicted in Fig. 4 and Fig. 5, 5 × 10^5^ CGNs were seeded onto glass bottom plates (3970-035, AGC Techno Glass Co., Ltd, Shizuoka, Japan) coated with poly-L-ornithine and affixed to a silicon chamber (flexiPERM mini; 94.6032.039, Sarstedt, Nümbrecht, Germany). In the experiments presented in Fig. 6, 7.5 × 10^5^ CGNs, introduced with plasmids as described below, were plated either on glass bottom plates or film bottom plates with grids (81166, Ibidi, Gräfelfing, Germany).

**Fig. 1.**
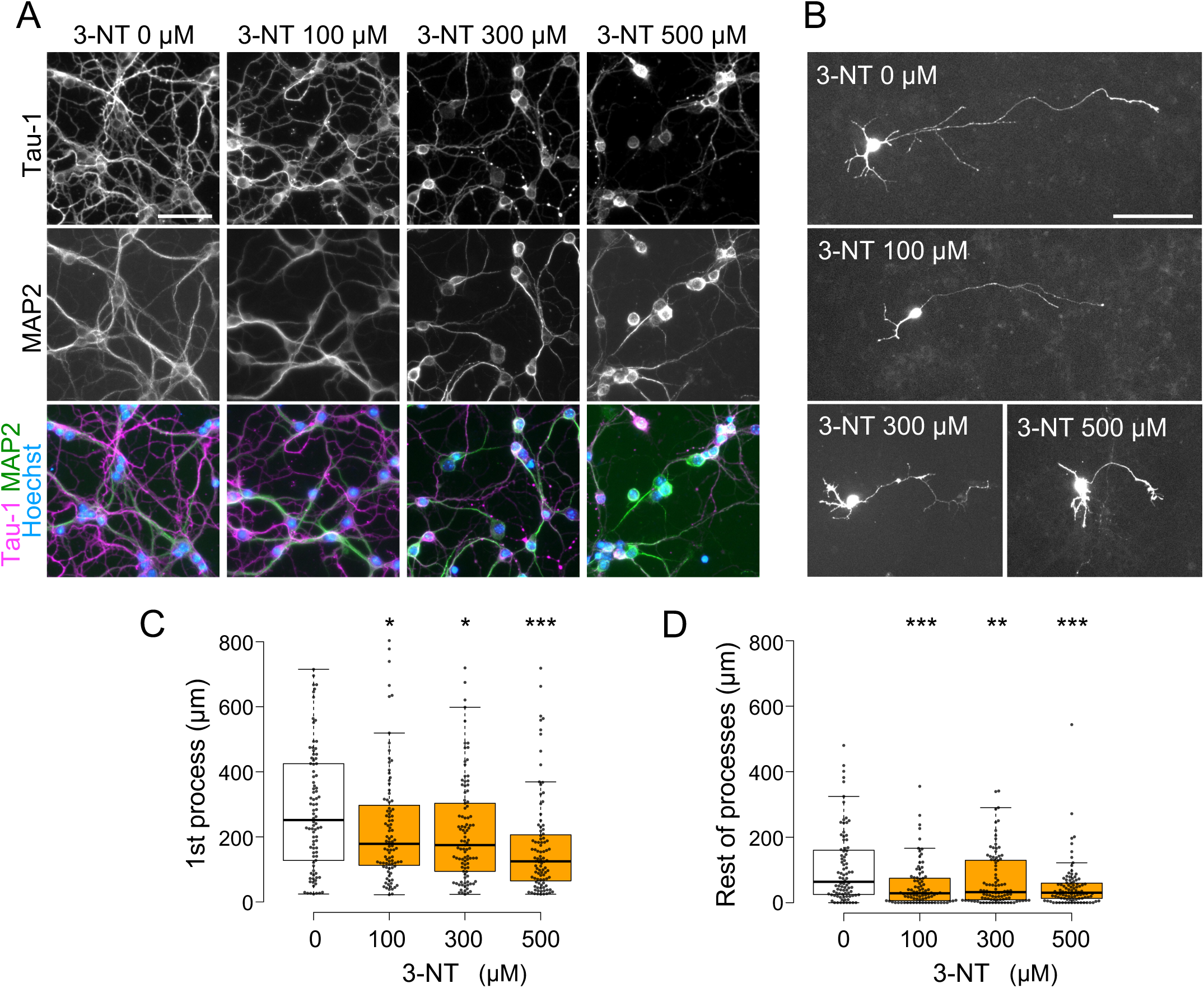
3-NT reduced the axonal length in CGNs. (A) CGNs at 4 days in vitro (DIV) were exposed to varying concentrations of 3-NT for 1 day, followed by immunocytochemical analysis utilizing Tau-1 (axonal marker; magenta) and MAP2 (dendritic marker; green) antibodies. Nuclear staining using Hoechst dye (blue) is revealed. (B) CGNs at 5 DIV, transduced with a vector expressing EGFP, were subjected to 3-NT treatment for 1 day. (C, D) Quantification of the length of the longest (axonal) process (C) and the rest of the processes (D) in each neuron was performed. Whiskers denote the range extending 1.5 times the interquartile range above and below the box limits. Data points are depicted as dots. 3-NT led to a reduction in the length of the longest process compared to the control condition. The Wilcoxon rank-sum test with the Benjamini-Hochberg correction was employed for comparison against 0 μM (white boxes) of 3-NT (n > 90 neurons from at least 3 experiments per condition). \**p* < 0.05, \*\**p* < 0.01, \*\*\**p* < 0.001. Neurons with processes less than 20 μm were excluded from analysis. Scale bars represent 50 μm.

**Fig. 2.**
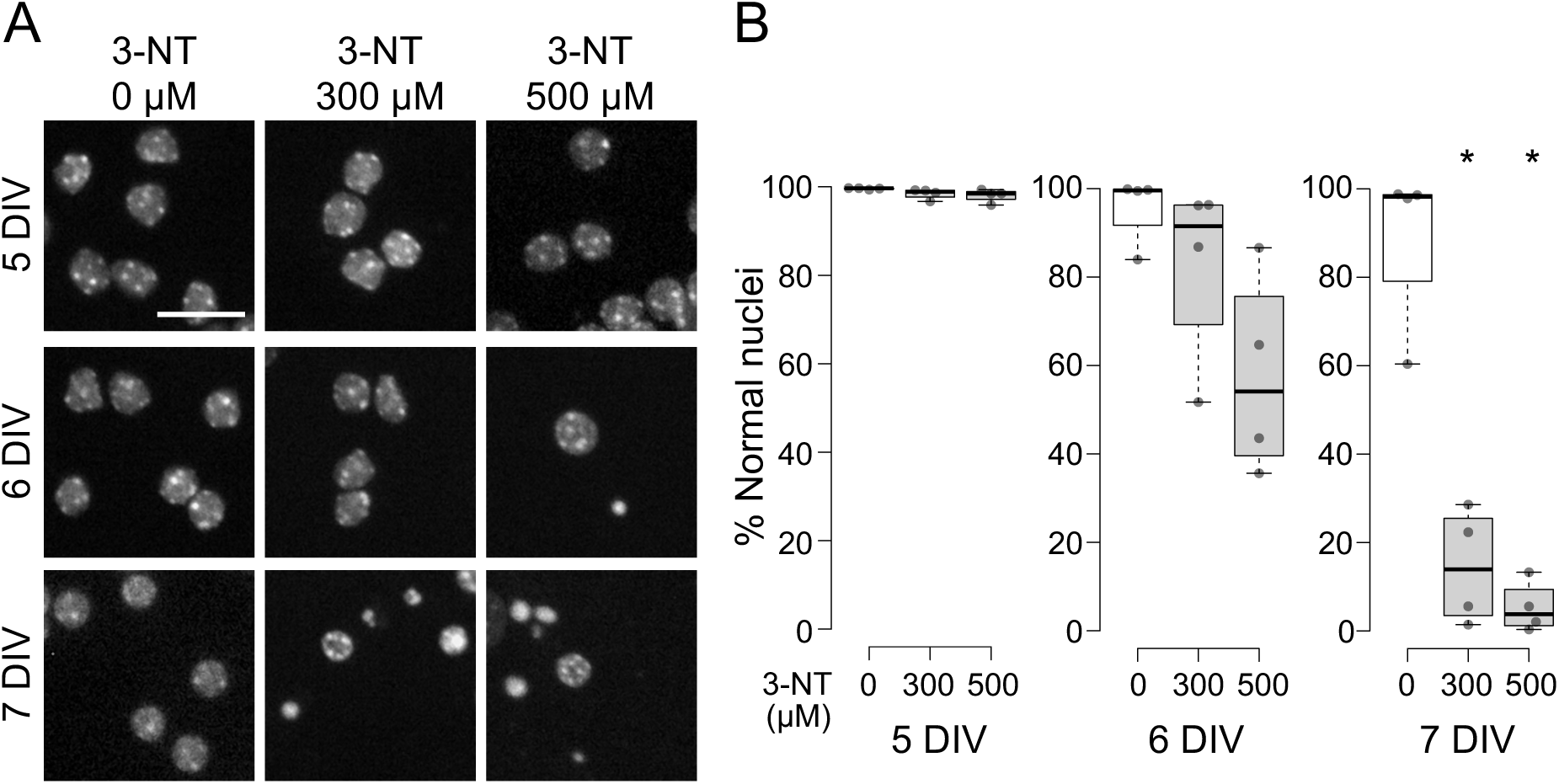
Effect of 3-NT on the cell survival of CGNs. (A) CGNs at 4 DIV were exposed to 0 μM, 300 μM, or 500 μM of 3-NT and fixed at 5 to 7 DIV (1 to 3 days post-treatment), followed by Hoechst nuclear staining. Treatment with 3-NT often resulted in neurons exhibiting condensed nuclei. (B) The percentage of neurons with normal nuclei was determined from the data in (A). Box plots were constructed as described in Fig. 1. No significant reduction in the percentage of normal nuclei was observed following 1-day treatment (5 DIV), whereas treatment with 300 μM or 500 μM of 3-NT for 2 and 3 days (6 and 7 DIV) significantly decreased the proportion of neurons with normal nuclei compared to 0 μM of 3-NT (white boxes). The Wilcoxon rank-sum test with the Benjamini-Hochberg correction was applied (n = 4 wells). \**p* < 0.05. The scale bar represents 20 μm.

**Fig. 3.**
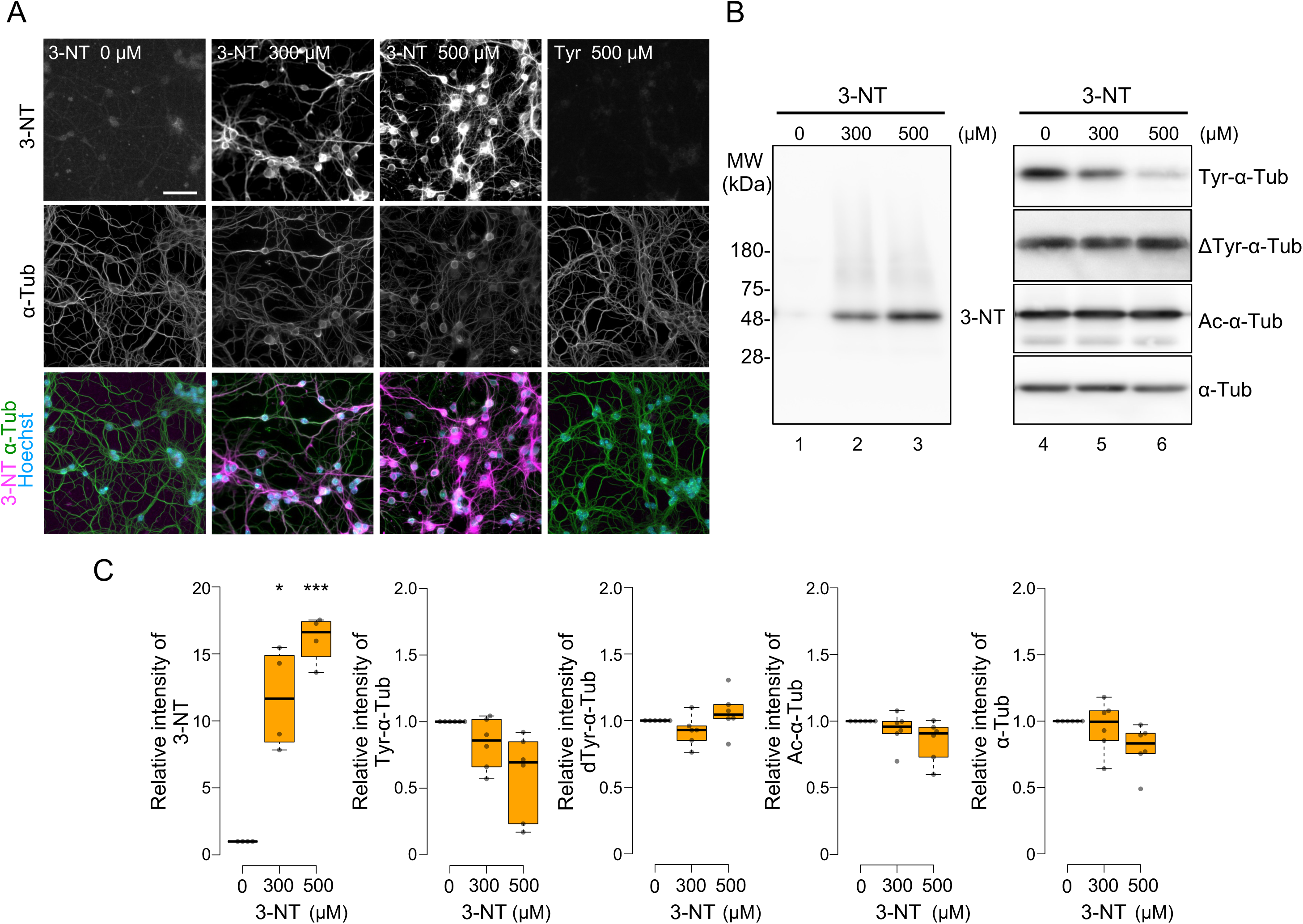
Selective incorporation of 3-NT into tubulins. (A) CGNs at 4 DIV were exposed to 0 μM, 300 μM, or 500 μM of 3-NT for 1 day, followed by immunocytochemistry using antibodies against 3-NT (magenta) and α-tubulin (green). Additionally, CGNs were treated with 500 μM tyrosine. Lysates of CGNs at 4 DIV, treated with different concentrations of 3-NT, were subjected to immunoblotting using antibodies against 3-NT (left panel) or antibodies against tyrosinated α-tubulin, detyrosinated α-tubulin, acetylated α-tubulin, or total α-tubulin regardless of its posttranslational modifications (right panel). (C) Quantification of signals from the immunoblotting depicted in (B) was performed. Treatment with 3-NT significantly enhanced the 3-NT signal corresponding to the mobility of α-tubulin, while it tended to decrease the signal for tyrosinated α-tubulin. Box plots were constructed as in Fig. 1. No significant changes were observed in signals for detyrosinated α-tubulin, acetylated α-tubulin, and total α-tubulin. The one sample *t*-test with the Benjamini-Hochberg correction was applied (n = 4 (3-NT antibody) or 6 (other antibodies)). \**p* < 0.05, \*\*\**p* < 0.001. The scale bar represents 50 μm.

**Fig. 4.**
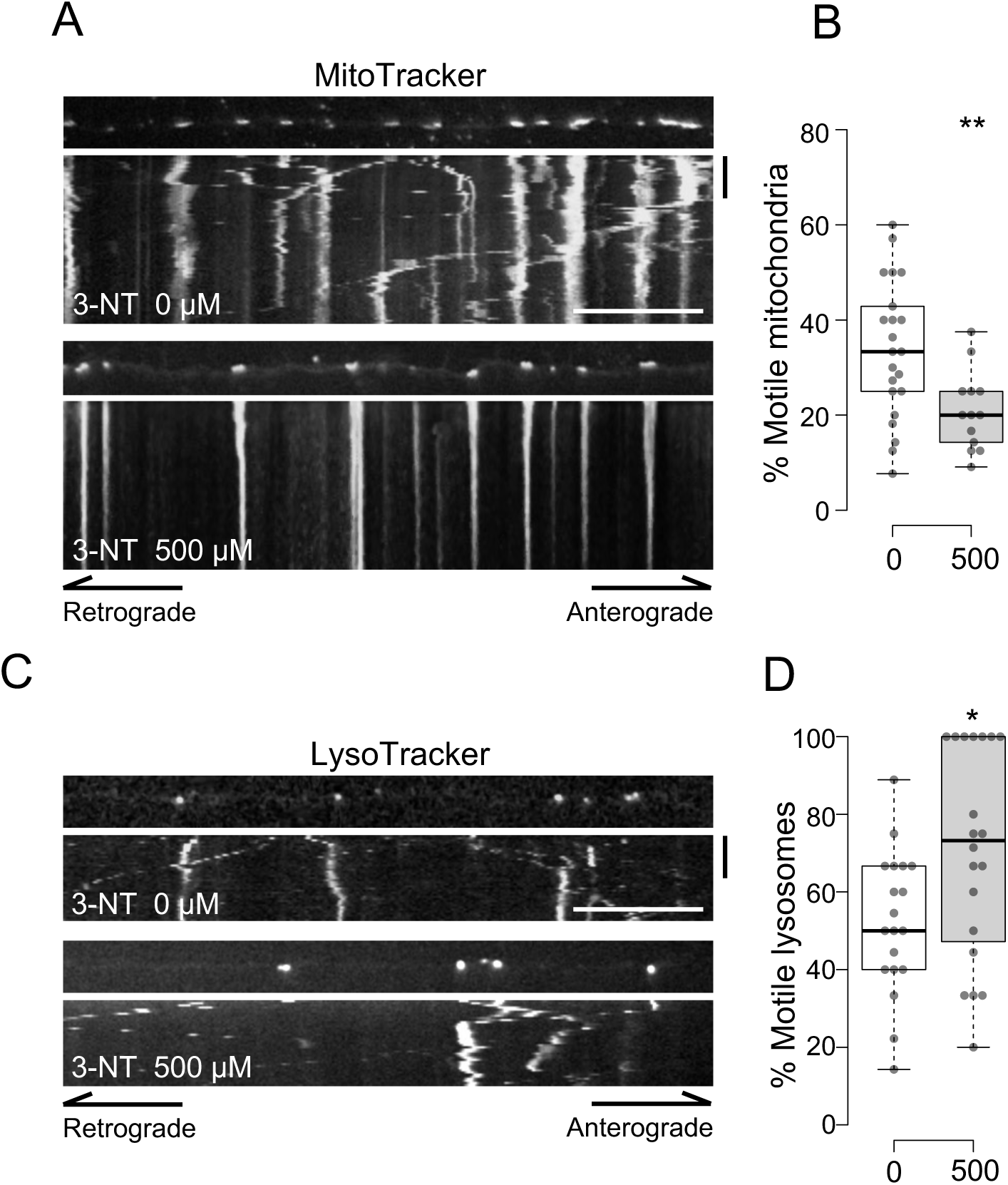
Inhibition of mitochondrial motility by 3-NT. (A, B) CGNs at 3 DIV, treated with (500 μM) or without (0 μM) 3-NT for 8 hours, were stained with MitoTracker and subjected to time-lapse imaging. Images were captured every 15 seconds for 20 minutes, and mitochondria exhibiting translocation of more than 5 μm were categorized as motile. 3-NT significantly decreased the ratio of motile mitochondria. Welch’s *t*-test was utilized (n = 22 (0 μM) and 13 (500 μM) axons from at least 4 experiments). (C, D) CGNs at 3 DIV, treated with or without 3-NT for 8 hours, were stained with LysoTracker and subjected to timelapse imaging for 10 minutes at 15-second intervals. 3-NT treatment increased the percentage of motile lysosomes. The Wilcoxon rank-sum test was employed (n = 19 (0 μM) and 20 (500 μM) axons from 4 experiments). Box plots were constructed as described in Fig. 1. \**p* < 0.05, \*\**p* < 0.01. Scale bars represent 20 μm and 5 seconds.

**Fig. 5.**
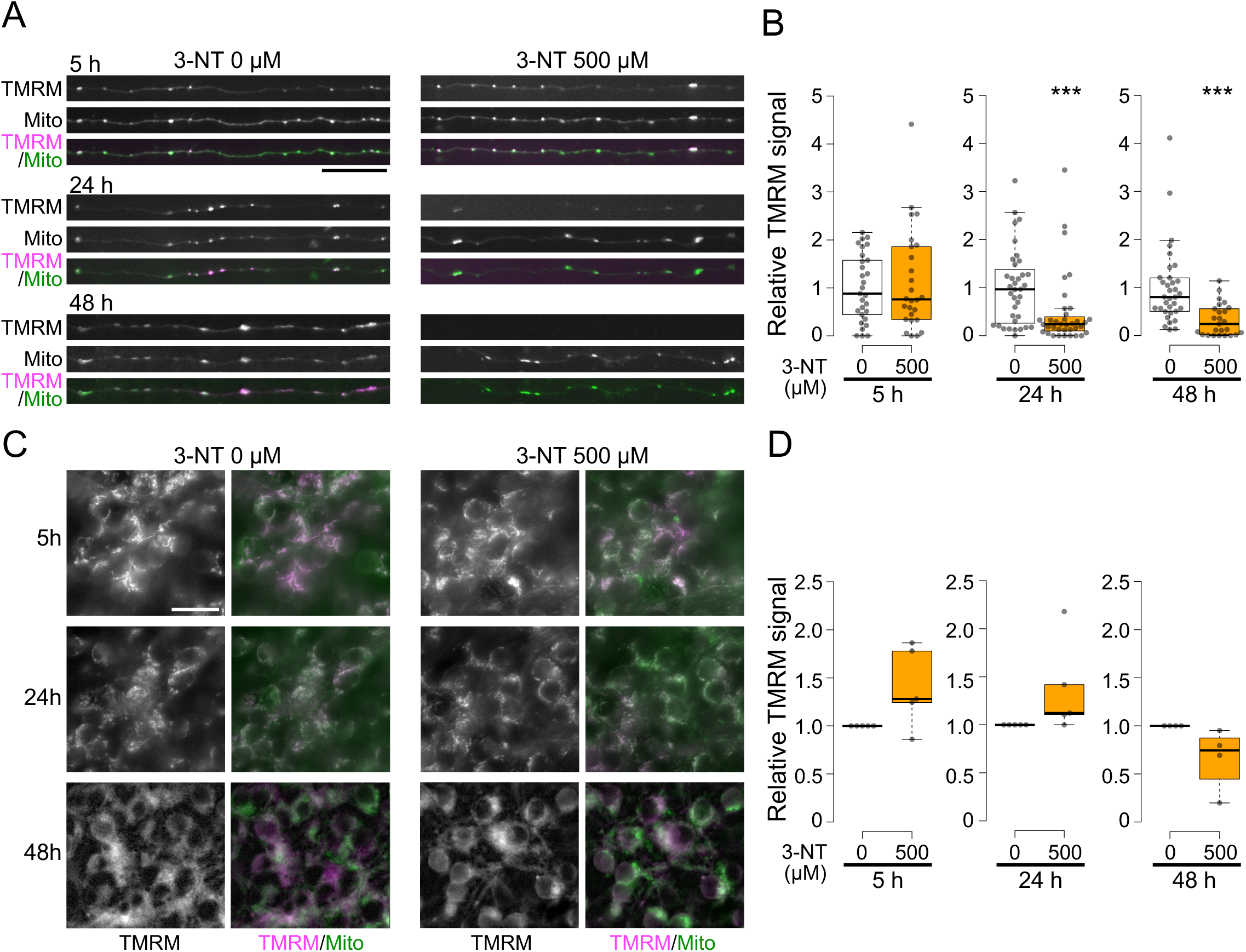
3-NT-induced loss of mitochondrial membrane potential in axons. CGNs at 4 DIV were exposed to 3-NT for 5, 24, or 48 hours and stained with TMRM (magenta) and MitoTracker (green). (A, B) Mitochondrial signals in axons were visualized (A), and signals in axonal segments (up to 100 from terminal ends) were quantified (B). In cultures treated with 3-NT, axonal mitochondria exhibited reduced TMRM signal after 24 and 48 hours of treatment. The Wilcoxon rank-sum test was employed (n > 24 axons from at least 5 experiments). (C, D) Images of TMRM and MitoTracker staining at the cell body location (C). TMRM signals at 500 μM of 3-NT were averaged and expressed relative to the signal at 0 μM of 3-NT. The Wilcoxon signed-rank test was utilized (n = 5 (5 h and 24 h) and 4 (48 h) experiments). Box plots were constructed as described in Fig. 1. \*\*\**p* < 0.001. The scale bar represents 20 μm.

**Fig. 6.**
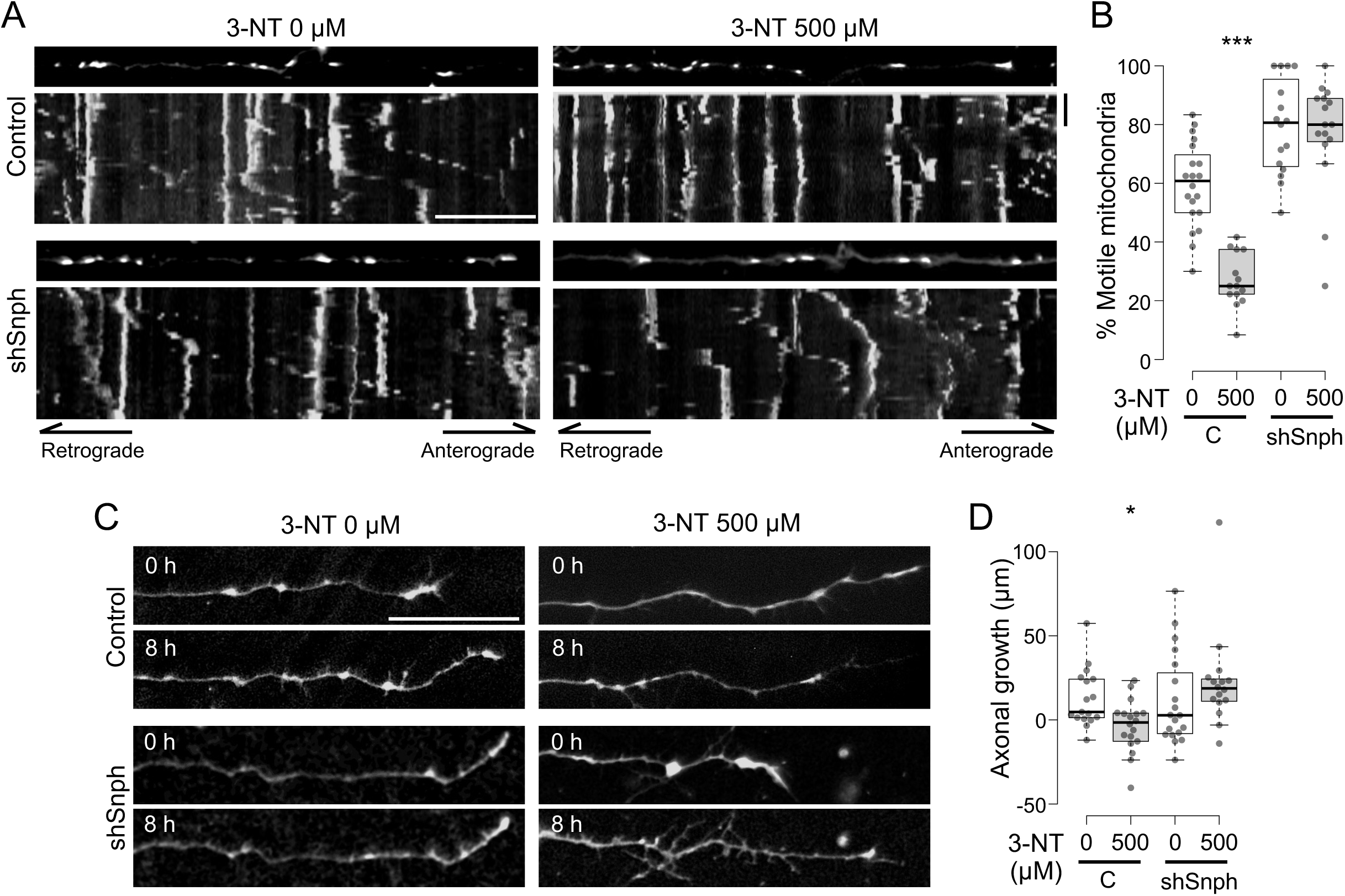
Rescue of 3-NT-induced inhibition of axonal growth by Snph inhibition. (A) Axons of CGNs at 5 DIV, transduced with the shSnph expression vector or its control along with the vector for tdTomato-Mito to visualize mitochondria, were treated with or without 3-NT for 8 hours, followed by time-lapse imaging for 20 minutes at 30-second intervals. (B) The percentage of motile mitochondria in each axon was determined as in Fig. 4B, and a comparison was made between 0 μM (white box) and 500 μM (gray box) of 3-NT. 3-NT reduced the percentage of motile axonal mitochondria in CGNs introduced with the control vector but not in those introduced with shSnph. (C) Axons of CGNs at 5 DIV, transduced with the shSnph expression vector or its control along with the vector for EGFP, were treated with or without 500 μM of 3-NT. Images were captured before (0 h) or after (8 h) the treatment. (D) The difference in axonal length before and after treatment was calculated. 3-NT inhibited axonal growth in CGNs introduced with the control vector but not in those introduced with shSnph. The Wilcoxon rank-sum test with the Benjamini-Hochberg correction was employed (n > 15 axons from at least 3 experiments). Box plots were constructed as described in Fig. 1. \**p* < 0.05, \*\*\**p* < 0.001. The scale bar represents 20 μm.

### 2.2. Gene transfer

Plasmids were introduced into CGNs using either the calcium phosphate method (Fig. 1B) or electroporation (Fig. 6). In the experiment shown in Fig. 1B, CGNs at 3 days in vitro (DIV) in 12-well plates were incubated in Dulbecco’s modified Eagle’s medium (DMEM, 045–30285, FUJIFILM Wako Pure Chemical Corporation) at 37°C in a CO_2_ chamber prior to transfection. A mixture containing 40 μl of 2 × HBS solution (270 mM NaCl, 9.5 mM KCl (191-01665, 163-03545, FUJIFILM Wako Pure Chemical Corporation), 1.4 mM NaH_2_PO_4_, 15 mM glucose (31718-15, 16806-15, Nacalai Tesque, Kyoto, Japan), 42 mM Hepes-KOH pH 7.1) and 40 μl of 250 mM CaCl_2_ (06731-05, Nacalai Tesque) solution, along with 0.4 μg of pmCherry-N1 (632523, Takara-bio, Shiga, Japan) and 0.4 μg of bcl-xl/pcDNA3 (Konishi et al., 2004), was prepared and added to the culture. After 15 minutes, the mixture was incubated with the neurons for an additional 15 minutes in a CO_2_ incubator. Subsequently, the neurons were washed twice with DMEM and placed in a conditioned medium. In the experiment shown in Fig. 6, 3 × 10^6^ CGNs were mixed with 2 μg of a Mission shRNA vector for Snph (TRCN0000201959, Merck, Darmstadt, Germany) or pLKO.1-TRC-control (Addgene_#10879) (Moffat et al., 2006), 2 μg of tdTomato-Mito-7 (Addgene_#58115) (Ai et al., 2008), 2 μg of pEGFP-C1 (Takara-bio, discontinued by supplier), and 1.5 μg of bcl-xl/pcDNA3 in 70 μl of DMEM. The CGNs were then placed in 2 mm cuvettes and subjected to decay pulses using a CUY21-edit II pulse generator (BEX, Tokyo, Japan) as follows: poring pulse, 275 V for 1 ms; driving pulse, +20 V for 50 ms repeated 5 times at 50 ms intervals, with reversal of polarity.

### 2.3. Immunocytochemistry

Immunocytochemistry of CGNs was performed as described previously (Seno et al., 2016; Imanaka et al., 2022). CGNs cultured in 12-well plates, treated with or without 3-NT (N0905) or tyrosine (T0550) (Tokyo Chemical Industry Co., Ltd., Tokyo, Japan), were fixed for 20 minutes at room temperature with 4% paraformaldehyde (163–18435, FUJIFILM Wako Pure Chemical Corporation) in phosphate-buffered saline (PBS), followed by treatment with 0.4% polyoxyethylene octylphenyl ether (169–21105, FUJIFILM Wako Pure Chemical Corporation) in PBS for an additional 20 minutes at room temperature. Afterward, the cells were incubated in a blocking solution containing 5% goat serum (16210–064, Thermo Fisher Scientific), 3% bovine serum albumin (BAC62, Equitech-Bio, Kerrville, TX), and 0.02% polyoxyethylene sorbitan monolaurate (166–21115, FUJIFILM Wako Pure Chemical Corporation) in PBS for 1 h at room temperature.

Following the blocking step, CGNs were treated with primary antibodies diluted in the blocking solution and incubated overnight at 4°C. Primary antibodies used included mouse monoclonal antibodies against Tau-1 (1:1000; AB5622, Merck) or α-tubulin (1:1000; 12G10, AB_1157911, Developmental Studies Hybridoma Bank, University of Iowa, Iowa City, IA), and rabbit polyclonal antibodies against MAP2 (1:1000; AB5622, Merck) or 3-NT (1:1000; 06-284, Merck). Following primary antibody incubation, CGNs were washed and incubated with secondary antibodies diluted in the blocking solution containing 1% goat serum for 2-3 hours at room temperature. Secondary antibodies used were goat anti-mouse IgG conjugated to Alexa Fluor 488/568 (1:1000; ab150113/ab175473, Abcam, Cambridge, UK) and goat anti-rabbit IgG conjugated to AlexaFluor 488/568 (1:1000; A11008/A11011, Thermo Fisher Scientific).

### 2.4. Western blotting

CGNs were washed with PBS and dissolved with boiled 2×SDS sample Buffer (125 mM Tris-HCl (207-06275, 084-05425, FUJIFILM Wako Pure Chemical Corporation) pH6.8, 4 % SDS (31606-75, Nacalai Tesque), 20 % Glycerol (075-00616, FUJIFILM Wako Pure Chemical Corporation) and heated for 10 minutes at 90 ℃. After a brief sonication the protein concentration was detected using a Pierce BCA kit (23227, Thermo Fisher Scientific), and heated for additional 2 minutes with 10 % of 2-mercaptoethanol and a small amount of bromophenol blue (137-06862, 021-02911, FUJIFILM Wako Pure Chemical Corporation). For the detection of 3-NT, no 2-mercaptoethanol was added. Equal amounts of protein were subjected to 5 % – 16 % semi-gradient SDS polyacrylamide gel electrophoresis, followed by transferring onto the FluoTrans W membrane (Pall Corporation, Port Washington, NY). After in incubating in the blocking solution (5 % goat serum, 25 mM Tris-HCl pH 7.5, 150 mM NaCl, 0.1 % polyoxyethylene sorbitan monolaurate) for 30 minutes at room temperature, the membranes were treated with primary antibodies in the blocking solution at 4 ℃ overnight, followed by incubation with secondary antibodies in the blocking solution with 1 % goat serum for 1 hour at room temperature. For primary antibodies, mouse monoclonal antibodies for tyrosinated tubulin (1:20000, clone TUB-1A2, T9028, Sigma-Aldrich), acetylated tubulin (1:10000, clone 6-11B-1, T6793, Sigma-Aldrich) or α-tubulin (1:1000; 12G10) and rabbit polyclonal antibodies against detyrosinated tubulin (1:10000, AB3021, Merck) and 3-NT (1:1000, 06-284, Merck) were used. For secondary antibodies, goat anti-mouse-IgG or anti-rabbit-IgG conjugated to horse radish peroxidase (1:2500, 330/458, Medical & Biological Laboratories Co., Ltd., Tokyo, Japan) were used.

### 2.5. Cell imaging and data analysis

Images were captured using an Axiovert 200 M microscope equipped with an Axiocam MRm digital camera controlled by AxioVision software (Carl Zeiss, Oberkochen, Germany). For live-cell imaging, CGNs were maintained at 36.7°C with 5% CO_2_ using a stage-top incubator (ZILCS, Tokaihit, Shizuoka, Japan). Image data were processed using ImageJ software (National Institute of Health, Bethesda, MD). The acquisition and processing of image data from fixed neurons (Fig. 1, 2) were conducted in a blinded manner. To analyze the process length of CGNs (Fig. 1), all neurites with a length of at least 20 μm emerging from the cell body were measured as described previously (Konishi et al., 2004). Nucleus integrity in Fig. 2 was determined by staining with the DNA dye Hoechst 33258 (861405, Merck) as described previously (Saga et al., 2020). To stain mitochondria, CGNs were washed with DMEM and treated with MitoTracker Red CM-H2XRos (500 nM, M7513), MitoTracker Green FM (200 nM, M7514) (Thermo Fisher Scientific), or tetramethylrhodamine methyl ester (TMRM) (250 nM, 203-1841, FUJIFILM Wako Pure Chemical Corporation) for 30 minutes in DMEM, then returned to conditioned media for imaging. Lysosomes were stained using LysoTracker Red DND-99 (500 nM, L7528, Thermo Fisher Scientific). Time-lapse imaging (Fig. 4, 6A) was conducted as described previously (Matsumoto et al., 2022). Kymograph data were quantified, and mitochondrial trajectories were analyzed based on the values, with mitochondria or lysosomes showing a maximum change in position during observation of more than 5 μm being considered motile. The axon for data analysis was predetermined from those with a length of at least 100 μm and fine mitochondrial (Fig. 4A, 6A), lysosomal (Fig. 4B), or EGFP (Fig. 6C) signals. In the experiments illustrated in Fig. 5, positions to acquire images were determined based on mitochondrial signals without observing TMRM signals. Statistical analyses were performed using R software (version 4.1.2). Normality was tested by the Shapiro-Wilk test. The Wilcoxon rank-sum test or the Wilcoxon signed-rank test was used for nonparametric analysis. The Welch’s *t*-test or the one-sample *t*-test was used for parametric analysis. The Benjamini-Hochberg correction is used to adjust for multiple comparisons. No calculation for sample size predetermination, no test for outliers, and no elimination of data points were performed. BoxplotR (http://shiny.chemgrid.org/boxplotr/) was used to generate boxplots. The levels of significance are denoted as follows: \**p* < 0.05, \*\**p* < 0.01, \*\*\**p* < 0.001. The values are presented as the median (interquartile range).

## 3. Results

### 3.1. 3-NT reduced the axonal length of CGNs

To investigate the potential impact of 3-NT on axonal morphology in non-DA neurons, dissociation cultures of CGNs were treated with varying concentrations of 3-NT. Previous research demonstrated that 3-NT induced cell death at concentrations of 250 to 500 μM in DA cells but not in non-DA cells (Blanchard-Fillion et al., 2006). In our study, CGNs were treated with different concentrations of 3-NT (i.e., 100 μM, 300 μM, and 500 μM) and subsequently subjected to immunocytochemical staining using markers for axons (Tau-1) and dendrites (MAP2). When CGNs were exposed to 3-NT before fixation at 5 DIV, we observed a tendency for reduction in axonal processes positive for Tau-1, particularly at higher concentrations (Fig. 1A). To quantitatively assess the effect on axons, CGNs were transfected with a vector for EGFP and then treated with varying concentrations of 3-NT. Under these conditions, the long, thin axonal processes, and short dendritic processes of individual CGNs could be visualized (Konishi et al., 2004) (Fig. 1B). The length of the longest (1st) process, representative of axonal length, as well as the rest of the processes of each neuron, were measured (Fig. 1B, C). Neurons with the 1st process length of less than 20 μm were excluded from the analysis to minimize contamination by dying cells or non-neuronal cells. Our findings revealed that 3-NT significantly reduced the length of the 1st process with just one day of treatment, even at 100 μM, and this effect on the length of the 1st process increased in a dose-dependent manner (257 (425–134) μm at 0 μM versus 125 (207–65) μm at 500 μM, *p* < 0.001, Wilcoxon rank-sum test). Moreover, the length of the rest of the processes was also reduced upon 3-NT treatment (64 (159–25) μm at 0 μM versus 30 (60–13) μm at 500 μM, *p* < 0.001, Wilcoxon rank-sum test), possibly due to the effect on the second axon and/or dendrites. Nevertheless, our results indicate that 3-NT shortens the axons of CGNs.

Next, we examined the effect of 3-NT on the survival of CGNs. CGNs at 4 DIV were exposed to 3-NT at concentrations of 300 μM and 500 μM for 1 to 3 days, followed by staining with Hoechst dye to visualize chromatin structure. In control CGN cultures, several heterochromatin foci were visible in almost all nuclei, whereas in cultures treated with 3-NT, condensed or fragmented nuclei were observed (Fig. 2A). There was no significant difference in the percentage of normal nuclei at one day after 3-NT treatment (5 DIV). However, by the second day after treatment (6 DIV), the percentage of normal nuclei began to decrease (100 (100–96) % at 0 μM versus 54 (70–42) % at 500 μM, *p* = 0.11, Wilcoxon rank-sum test), and by the third day after treatment (7 DIV), most cells exhibited abnormal nuclei (98 (99–88) % at 0 μM versus 4 (7–2) % at 500 μM, *p* < 0.05, Wilcoxon rank-sum test) (Fig. 2 A, B). These results suggest that 3-NT treatment could lead to neuronal toxicity in CGNs. Considering that 3-NT shortened axonal length within one day (Fig. 1), it is likely to be the primary effect rather than a consequence of neuronal death.

### 3.2. 3-NT is preferentially incorporated in α-tubulins in CGNs

To elucidate the mechanism by which 3-NT shortens the axonal length of CGNs, we investigated the localization of 3-NT incorporation in our system. CGNs were exposed to 3-NT at concentrations of 300 μM and 500 μM for one day and subjected to immunocytochemistry using an antibody against 3-NT. Signals for 3-NT were detected not only in the cytoplasmic region but also in neuronal processes exhibiting tubulin signals in CGNs exposed to 3-NT, whereas no signals were observed in control neurons without treatment or those treated with tyrosine (Fig. 3A). Western blotting of CGN lysate revealed that most of the 3-NT signals were detected at the position corresponding to tubulins and were increased by more than 10-fold with 3-NT treatment (16.6 (17.3–15.4) fold at 500 μM, *p* < 0.001 relative to 0 μM, one-sample *t*-test) (Fig. 3B, C). These observations are in accordance with the notion that free 3-NT is incorporated in the carboxy-terminal end of α-tubulin instead of tubulin by Ttl (Eiserich et al., 1999). Indeed, the signal for tyrosination at the carboxy-terminal end of α-tubulin tended to be reduced by 3-NT treatment (0.69 (0.82–0.34) fold at 500 μM, *p* = 0.051 relative to 0 μM, one-sample *t*-test) (Fig. 3B, C). No significant changes were observed in signals for detyrosinated α-tubulin, acetylated α-tubulin, or the amount of α-tubulin (Fig. 3B, C). These results indicate that 3-NT is incorporated into α-tubulin within neurites in correlation with axonal shortening in CGNs.

### 3.3. 3-NT inhibited mitochondrial motility

Previous studies have reported that tyrosination/detyrosination of α-tubulins regulates kinesin-1-mediated anterograde axonal transport as well as dynein-mediated retrograde axonal transport (Konishi and Setou, 2009; McKenney et al., 2016; Nirschl et al., 2016). Since mitochondria are transported by kinesin-1/dynein and play crucial roles in the growth and maintenance of axons (Hirokawa et al., 2010; Saxton and Hollenbeck 2012; Sheng and Cai 2012; Smith and Gallo 2018; Cheng et al., 2022), we investigated whether 3-NT affects mitochondrial transport in CGN axons. By labeling with MitoTracker, both motile and stationary mitochondria were observed in axons of control CGNs at 3 DIV (Fig. 4A), consistent with the previous report (Matsumoto et al., 2022). In CGN axons treated with 3-NT for 8 hours, mitochondria tended to remain stationary during observation (Fig. 4A). Mitochondrial positions were measured from the kymograph, and mitochondria with a maximum change in position of more than 5 μm during a 20-minute observation were considered motile. The proportion of motile mitochondria significantly decreased following 3-NT treatment (33.3 (42.1–25.0) % at 0 μM versus 20.0 (25.0–14.3) % at 500 μM, *p* < 0.01, Welch’s *t*-test) (Fig. 4B). We also monitored the axonal transport of lysosomes using LysoTracker, which stains acidic granules, as kinesin-1 and dynein have been reported to be involved in the axonal transport of lysosomes (Roney et al., 2022) (Fig. 4C). This experiment was conducted for 10 minutes, as the motility of acidic granules was greater than that of mitochondria. In contrast to mitochondrial transport, the motility of acidic granules in CGN axons was not reduced by exposure to 3-NT, but rather significantly increased (50.0 (66.7–40.0) % at 0 μM versus 73.2 (100.0–48.6) % at 500 μM, *p* < 0.05, Welch’s *t*-test) (Fig. 4C D).

To further investigate the effect of 3-NT on mitochondrial function, CGNs were stained with TMRM, a fluorescent probe for mitochondrial membrane potential (MMP) (Buckman and Reynolds, 2001). At 5 hours post-treatment, the majority of axonal mitochondria were TMRM-positive, with no significant reduction observed following exposure to 3-NT (Fig. 5A, B). However, at 24 and 48 hours post-treatment, TMRM signals in axons were markedly reduced by 3-NT (24 h: 0.97 (1.33–0.28) at 0 μM versus 0.24 (0.37–0.11) at 500 μM, *p* < 0.001; 48 h: 0.80 (1.18–0.52) at 0 μM versus 0.24 (0.56–0.20) at 500 μM, *p* < 0.001, Wilcoxon rank-sum test). Conversely, in the cell body, no decrease in TMRM signal due to 3-NT was detected until 24 hours but rather tended to increase (1.12 (1.42–1.11) at 500 μM relative to 0 μM at 24 h, *p* = 0.06, Wilcoxon signed-rank test) (Fig. 5C, D). After 48 hours, the TMRM signal tended to decrease with 3-NT exposure, although it did not reach statistical significance (0.74 (0.83–0.57) at 500 μM relative to 0 μM, *p* = 0.06, Wilcoxon signed-rank test). These results suggest that the reduction of MMP in axons precedes that in the soma following 3-NT treatment.

### 3.4. 3-NT-mediated reduction of mitochondrial motility requires syntaphilin

Previous studies have reported that syntaphilin (Snph) plays a critical role in immobilizing axonal mitochondria by anchoring them to microtubules, and inhibition of Snph in neurons increases mitochondrial motility in axons (Kang et al., 2008; Sheng and Cai 2012; Courchet et al., 2013). Therefore, we investigated whether inhibiting Snph could enhance axonal mitochondrial motility, which was reduced by 3-NT. To address this question, CGNs were transfected with a vector expressing small hairpin RNA for Snph (shSnph) along with a vector expressing a mitochondria-targeting sequence tagged with tdTomato (tdTomato-Mito). In CGNs transfected with a non-hairpin control, treatment with 3-NT for 8 hours significantly decreased the motile mitochondria in axons (60.8 (68.2 – 50.0) % at 0 μM versus 25.0 (35.5 – 22.2) % at 500 μM, *p* < 0.001, Wilcoxon rank-sum test) (Fig. 6A, B). In CGNs expressing shSnph, the number of motile mitochondria in axons tended to be higher, consistent with previous studies (Kang et al., 2008; Courchet et al., 2013), and 3-NT did not cause a significant reduction in mitochondrial motility (80.6 (93.2 – 66.2) % at 0 μM versus 80.0 (88.9 – 74.6) % at 500 μM, *p* = 0.993, Wilcoxon rank-sum test) (Fig. 6A, B). These results indicate that 3-NT-dependent mitochondrial immobilization can be restored by inhibiting Snph.

### 3.5. Inhibition of SNPH rescues axonal growth in 3-NT treated CGNs

Given that inhibition of Snph reversed the effect of 3-NT on the reduction of mitochondrial motility, we tested whether it could also rescue the 3-NT-mediated axonal shortening. We focused on the short-term effects (i.e., within 8 h), as long-term treatment with 3-NT caused mitochondrial dysfunction and cell death. CGNs were transfected with a vector for shSnph together with a vector for EGFP to visualize the axonal morphology of transfected neurons. By comparing pre- and post- (8 h) 3-NT treatment, differences in the length of axons were quantified. Without treatment with 3-NT, both growing and contracting axons were detected, but an overall tendency to slightly elongate was observed (Fig. 6C, D). 3-NT treatment significantly reduced the growth of axons in control neurons, with more axons being retracted (4.7 (24.2 – 1.3) μm in 0 μM versus −1.6 (3.9 – −12.0) μm in 500 μM, *p* < 0.05, Wilcoxon rank-sum test) (Fig. 6C, D), reflecting the axonal shortening effect of 3-NT observed in Fig. 1 at least in part. However, in shSnph-expressing neurons, 3-NT did not cause a reduction in axonal growth. Overall growth even tended to increase with 3-NT in the shSnph expressing CGNs, although not statistically significant (2.7 (28.0 – −8.2) μm in 0 μM versus 18.7 (23.9 – 11.4) μm in 500 μM, *p* = 0.123, Wilcoxon rank-sum test) (Fig. 6C, D). These results indicate that increasing mitochondrial motility by knocking down Snph could rescue the effect of 3-NT to inhibit axonal growth. Collectively, these observations suggest that 3-NT causes axonal shortening in CGNs, which is mediated at least in part by immobilizing axonal mitochondria.

## 4. Discussion

### 4.1. The impact of of 3-NT on CGNs

The ramifications of 3-NT on neuronal structure have remained undetermined. In this study, we have revealed that 3-NT inhibits axonal growth in cultured CGNs (Fig. 1). Previous research has shown that free 3-NT is incorporated into the carboxy-terminal of α-tubulin, inducing alterations in cellular morphology and microtubule organization in non-neuronal cells (Eiserich et al., 1999; Bisig et al., 2002). Consistently, no clear bands for 3-NT other than tubulin were detected in Western blotting, suggesting that free 3-NT is selectively incorporated into α-tubulin (Fig. 3). Furthermore, the incorporation of 3-NT into α-tubulin was correlated with axonal shortening. The effect of 3-NT on axons began to appear from 100 μM, a concentration comparable to that detected in striatal pathological conditions (Pennathur et al., 1999). It is also possible that 3-NT acts over a longer period or locally, affecting axonal function *in vivo*.

In the current study, we observed that 3-NT treatment for 3 days induce cell death in CGN culture (Fig. 2). Previous studies suggested that 3-NT-mediated death of DA neurons is not related to α-tubulin modification, as 3-NT did not cause cell death in non-DA neurons under the same conditions, even though it incorporated 3-NT into α-tubulins. Additionally, an inhibitor for aromatic amino acid decarboxylase prevented the death of DA neurons triggered by 3-NT (Blanchard-Fillion et al., 2006). Apart from DA neurons, a report suggests that 3-NT induces apoptosis in motor neurons without incorporating into α-tubulin (Peluffo et al., 2004). Therefore, 3-NT-induced cell death in CGNs might occur through a pathway independent of incorporation into α-tubulin. On the other hand, the decrease in MMP in the axon followed by the decrease in the soma correlated well with cell death (Fig. 2, 5), may contribute, at least in part, to 3-NT-mediated cell death. In any case, the inhibitory effect of 3-NT on axonal outgrowth was detected before cell death and were rescued by increasing mitochondrial motility by knocking down Snph, suggesting that it is not merely a consequence of activation of cell death signaling.

### 4.2. The effect of 3-NT on the axonal transport machinery

Transport of mitochondria is mediated by kinesin-1 and dynein, which are anterograde and retrograde microtubule motors, respectively. Previous reports have revealed that these motors are regulated by tyrosination/detyrosination of α-tubulin (Konishi and Setou, 2009; McKenney et al., 2016; Nirschl et al., 2016). Thus, the possible mechanism for inhibiting mitochondrial motility by 3-NT incorporation into α-tubulins is the disruption of tyrosination/detyrosination-mediated regulation of kinesin-1/dynein motors. Accordingly, a previous study suggested that 3-NT reattributes cytoplasmic dynein in A549 epithelial cells by altering microtubule association (Eiserich et al., 1999). A more recent study revealed that DA neurons with a mutation in α-synuclein, exposed to agrochemicals, resulted in the inhibition of anterograde mitochondrial transport, thought to be mediated by the nitration of α-tubulins inhibiting the association of Kif5B, the heavy chain of kinesin-1, to microtubules (Stykel et al., 2018).

In the present study using CGNs, 3-NT-dependent inhibition of mitochondrial motility was rescued by Snph knockdown (Fig. 6A), and axonal transport of acidic lysosomes which is mediated by kinesin-1/3 and dynein (Roney et al., 2022) was not inhibited by 3-NT (Fig. 4C, D). These results suggest that the function of motors on microtubules was not completely blocked. Thus, it is possible that selective inhibition may occur due to the types of motors mentioned above, as well as adapter factors and other microtubule-related factors including Snph. While mitochondrial transport in the axon is crucial for its elongation and maintenance, inhibition of mitochondrial anchoring by knockdown of Snph results in the shortening of axonal branches in cortical neurons (Courchet et al., 2013; Zhou et al., 2016), suggesting that the balance of motile and stationary mitochondria is important for proper axonal morphogenesis. Consistently, 3-NT inhibited axonal growth and increased the proportion of retracted axons, whereas in neurons with Snph knockdown, even if it may have any promoting axonal outgrowth, no inhibitory effect was observed (Fig. 6C, D). This suggests that, at least in the short term, axonal growth can be restored through an appropriate balance of mitochondrial movement and arrest, even in the presence of 3-NT.

### 4.3 Conclusions

Our findings demonstrate that 3-NT inhibits axon growth and the motility of mitochondria in CGNs. Additionally, Snph knockdown reverses these effects induced by 3-NT. This study offers new evidence for the potential role of 3-NT and its mechanism in axonal dysfunction.

## Ethical declaration

Animal handling procedures adhered to institutional regulations for animal research and were approved by the Animal Ethical Committee of the University of Fukui (approval numbers: R01082, R02901, R03093).

## Author contributions

**Masahiro Hirai:** Investigation, Formal analysis. **Kohei Suzuki:** Investigation, Formal analysis. **Yusuke Kassai:** Investigation. **Yoshiyuki Konishi:** Conceptualization, Investigation, Formal analysis, Writing, Supervision.

## Data availability

Data will be made available on request.

## Acknowledgements

The pLKO.1-TRC-control was a gift from David Root. The tdTomato-Mito-7 was a gift from Dr. Michael Davidson.

## Abbreviations

CGN: cerebellar granule neuron
DA: dopaminergic
DIV: days *in vitro*
DMEM: dulbecco’s modified eagle’s medium
MMP: mitochondrial membrane potential
3-NT: 3-nitrotyrosin
PBS: phosphate-buffered saline
SDS: sodium dodecyl sulfate
Snph: syntaphilin
TMRM: tetramethylrhodamine methyl ester
Ttl: tubulin tyrosine ligase

